# Specific cardiolipin-SecY interactions are required for proton-motive-force stimulation of protein secretion

**DOI:** 10.1101/202184

**Authors:** Robin A. Corey, Euan Pyle, William J. Allen, Marina Casiraghi, Bruno Miroux, Ignacio Arechaga, Argyris Politis, Ian Collinson

## Abstract

The transport of proteins across or into membranes is a vital biological process, achieved in every cell by the conserved Sec machinery. In bacteria, SecYEG combines with the SecA motor protein for secretion of pre-proteins across the plasma membrane, powered by ATP hydrolysis and the trans-membrane proton-motive-force (PMF). The activities of SecYEG and SecA are modulated by membrane lipids, particularly by cardiolipin, a specialised phospholipid known to associate with a range of energy-transducing machines. Here, we identify two specific cardiolipin binding sites on the *Thermotoga maritima* SecA-SecYEG complex, through application of coarse-grained molecular dynamics simulations. We validate the computational data and demonstrate the conserved nature of the binding sites using *in vitro* mutagenesis, native mass spectrometry and biochemical analysis of *Escherichia coli* SecYEG. The results show that the two sites account for the preponderance of functional cardiolipin binding to SecYEG, and mediate its roles in ATPase and protein transport activity. In addition, we demonstrate an important role for cardiolipin in the conferral of PMF-stimulation of protein transport. The apparent transient nature of the CL interaction might facilitate proton exchange with the Sec machinery and thereby stimulate protein transport, by an as yet unknown mechanism. This study demonstrates the power of coupling the high predictive ability of coarse-grained simulation with experimental analyses, towards investigation of both the nature and functional implications of protein-lipid interactions.

**Significance Statement:** Many proteins are located in lipid membranes surrounding cells and cellular organelles. The membrane can impart important structural and functional effects on the protein, making understanding of this interaction critical. Here, we apply computational simulation to the identification of conserved lipid binding sites on an important highly conserved bacterial membrane protein, the Sec translocase (SecA-SecYEG), which uses ATP and the proton motive force (PMF) to secrete proteins across the bacterial plasma membrane. We experimentally validate and reveal the conserved nature of these binding sites, and use functional analyses to investigate the biological significance of this interaction. We demonstrate that these interactions are specific, transient, and critical for both ATP- and PMF- driven protein secretion.

## Introduction

The translocation of proteins across and into membranes is an essential cellular process, acting as a key step in the biogenesis of a diverse array of proteins. In every cell, a major portion of this process is handled by the structurally-conserved Sec translocon – SecYEG in prokaryotes and Sec61 in eukaryotes. At the bacterial plasma membrane, SecYEG can carry out translocation either post-translationally, using the SecA motor protein (Figure 1A), or co-translationally, interacting directly with the ribosome-nascent-chain complex.

**Figure 1.**
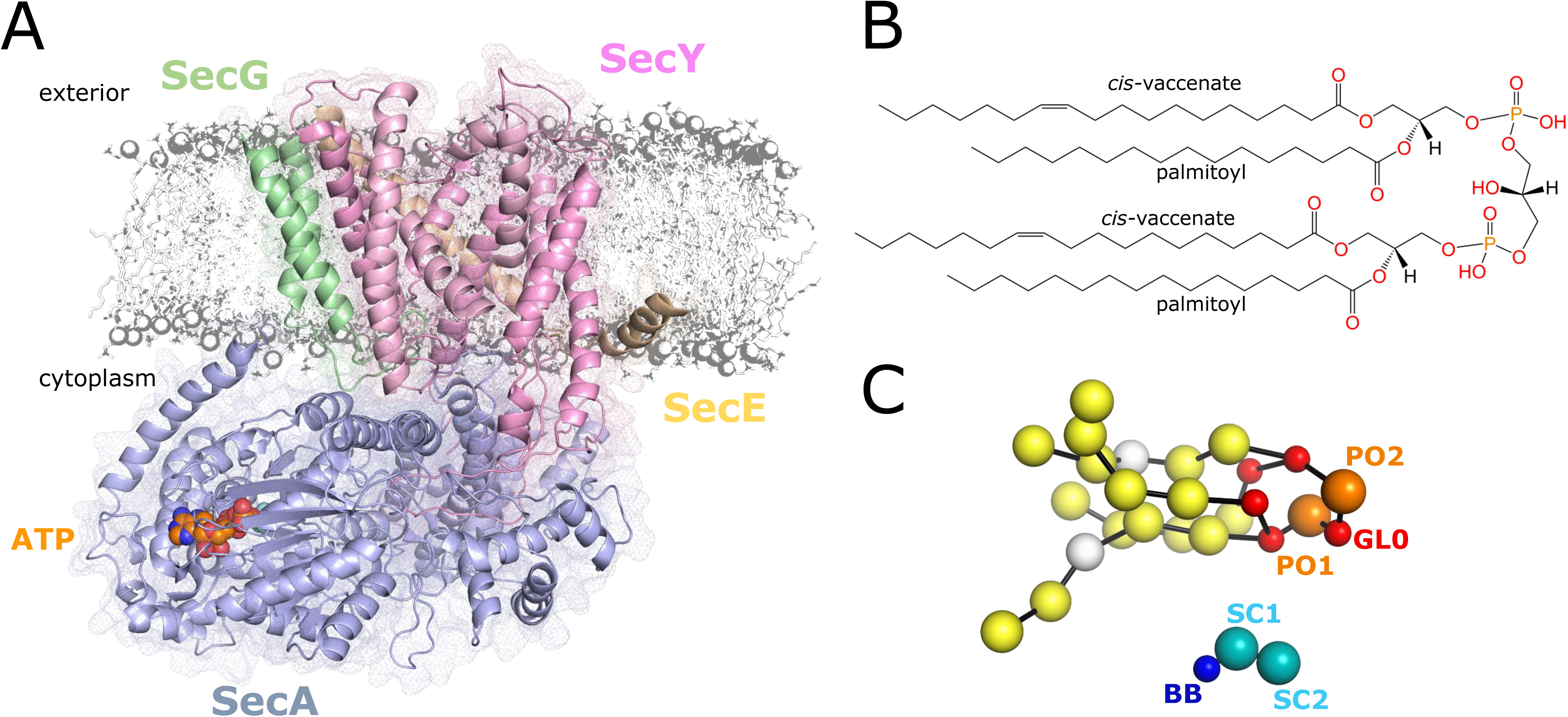
SecA-SecYEG structure and the MARTINI force field. A) Cartoon of SecYEG bound to SecA, from PDB 3DIN (25). The Sec subunits are shown as cartoon with mesh overlay, with SecY in light pink, SecE orange, SecG green and SecA light-blue. The ATP analogue (ADP-BeFx) is coloured as orange, blue and red spheres. A small region of lipid bilayer is represented in grey. B) Chemical structure of a common cardiolipin (CL) molecule, in this case di-1-palmitoyl-2-*cis*-vaccenate-phosphatidylglycerol. CL are defined by having four acyl tails and two phosphate head groups. Structure made using LipidMAPS online structure drawing tool (68). C) MARTINI representation of the CL molecule in panel B, with phosphate beads in orange, glycerol beads in red, and acyl tail beads in yellow and white (for beads containing a double bond). Below, a lysine molecule with side chain beads in light blue and the backbone bead (BB) in dark blue. In MARTINI, each bead represents approximately four heavy atoms plus associated hydrogens. Bonds connecting the beads are shown as black lines.

The post-translational pathway is primarily involved in protein secretion, providing the trafficking route for the bulk of periplasmic and outer membrane proteins, as well ass those proteins destined for the external medium. Pre-proteins, which contain a cleavable N-terminal signal sequence, are passed through a pore in the centre of SecY (1, 2), in a process powered by both ATP and the trans-membrane proton-motive-force (PMF) (3). How ATP binding and hydrolysis drives translocation is currently under debate (4-6), whereas the mechanism of PMF powered transport is completely unknown.

Like many integral membrane proteins (7, 8), SecYEG is functionally modulated by interactions with specific lipids. Our understanding of the nature of these interactions is limited due to the requirement of extracting proteins from the membrane, usually with detergents, for purification and characterisation; a treatment that removes some or all of the natively bound lipids. Various functional studies have aimed at addressing this, revealing that anionic phospholipids, *e.g.* cardiolipin (CL) and phosphatidylglycerol (PG), can stimulate SecA ATPase activity both alone and in complex with SecYEG (9, 10) and, moreover, are important for normal levels of translocation (11, 12). In addition, CL binding has been demonstrated to promote SecYEG dimer formation, between adjacent SecE transmembrane helices (10).

SecYEG is not unusual in its interaction with CL, which co-isolates with a considerable number of prokaryotic and mitochondrial energy-transducing membrane protein complexes (13). CL has a distinctive structure, comprising two PG molecules joined by a glycerol head group, giving it two phosphate head groups and four acyl chains (Figure 1B). It has been proposed that each phosphate on CL has a separate *pK_a_* (14), resulting in a bicyclic resonance structure at pH 7. This would allow CL to act as a reservoir for protons and buffer against localised shifts in pH (15), important for biological systems involving proton transfer. However, more recent analyses support similar *pK_a_* values for both phosphates (16, 17), meaning CL would carry a −2 charge at pH 7. How this would affect its ability to shuttle protons is uncertain.

CL binding sites on proteins are often typified by the presence of one or more basic residues that form pockets of positive charge on the surface of the protein (8, 13). This has been observed in structural data (18, 19), as well as in simulation data (20, 21) gathered using the MARTINI coarse-grained molecular dynamics (CGMD) force field (22, 23). Although of lower resolution than atomistic modelling, CGMD is particularly well suited to the study of protein-lipid interactions (24), primarily as a reduced particle count and longer step size produces significant improvement in computational efficiency. This permits both longer and larger systems for the same computational resource: a notable advantage when attempting to simulate the free diffusion of lipids within systems containing both bilayers and large multisubunit protein complexes.

Here, we adapt and extend these analyses towards identification of CL binding sites in *Thermotoga maritima* SecA-SecYEG (25). The data reveals the presence of two distinct CL binding sites in SecY, as well as supporting a previously reported (5, 26) lipid binding site on the N-terminus of SecA. We then validate the putative SecY CL binding sites and demonstrate the conserved nature of the SecY-CL interaction using biochemical and native mass spectrometry analyses of *Escherichia coli* SecYEG variants. In addition, we establish the importance of specific CL binding at these sites to heightened SecA-SecYEG activity, and demonstrate an important role in CL binding in the stimulation of translocation by the PMF. Finally, we demonstrate that whilst CL does mediate SecYEG dimer formation, this is probably *via* additional non-specific interactions.

## Results

### Coarse-grained analyses of Sec CL interaction using self-assembling membranes

We applied the MARTINI CG force field (22, 23) to predict Sec-CL interactions, using the crystal structure of SecA-SecYEG in an nucleotide-free state as an input model (Figure 1A) (25). In MARTINI, approximately 4 heavy atoms (*i.e.* C, P, O, N *etc.*) and associated hydrogens are modelled by a single bead, allowing CL (Figure 1B) to be represented by 25 beads (Figure 1C), including two phosphate head group beads (‘PO1’ and ‘PO2’) and a linking glycerol bead (‘GL0’). Amino acids are represented with a single bead for the backbone (‘BB’) and up to 4 beads for the side chain (‘SC1’, ‘SC2’, ‘SC3’ and ‘SC4’) (Figure 1C; blue beads).

Our initial analyses followed a ‘self-assembly’ approach, whereby free lipids (with CL at ~10%, as per a typical bacterial plasma membrane (27, 28)) were allowed to form membranes around a positionally-restrained protein over 1 μs simulations. Bilayer formation was verified using visual analysis with VMD (29), and the post-bilayer time frames from 10 independent simulations were combined into one trajectory. The occupancy of the CL head groups along the plane of the membrane was then computed using density analysis in VMD, with the data revealing key regions of increased CL density (Figure 2A), suggesting specific Sec-CL interactions.

**Figure 2.**
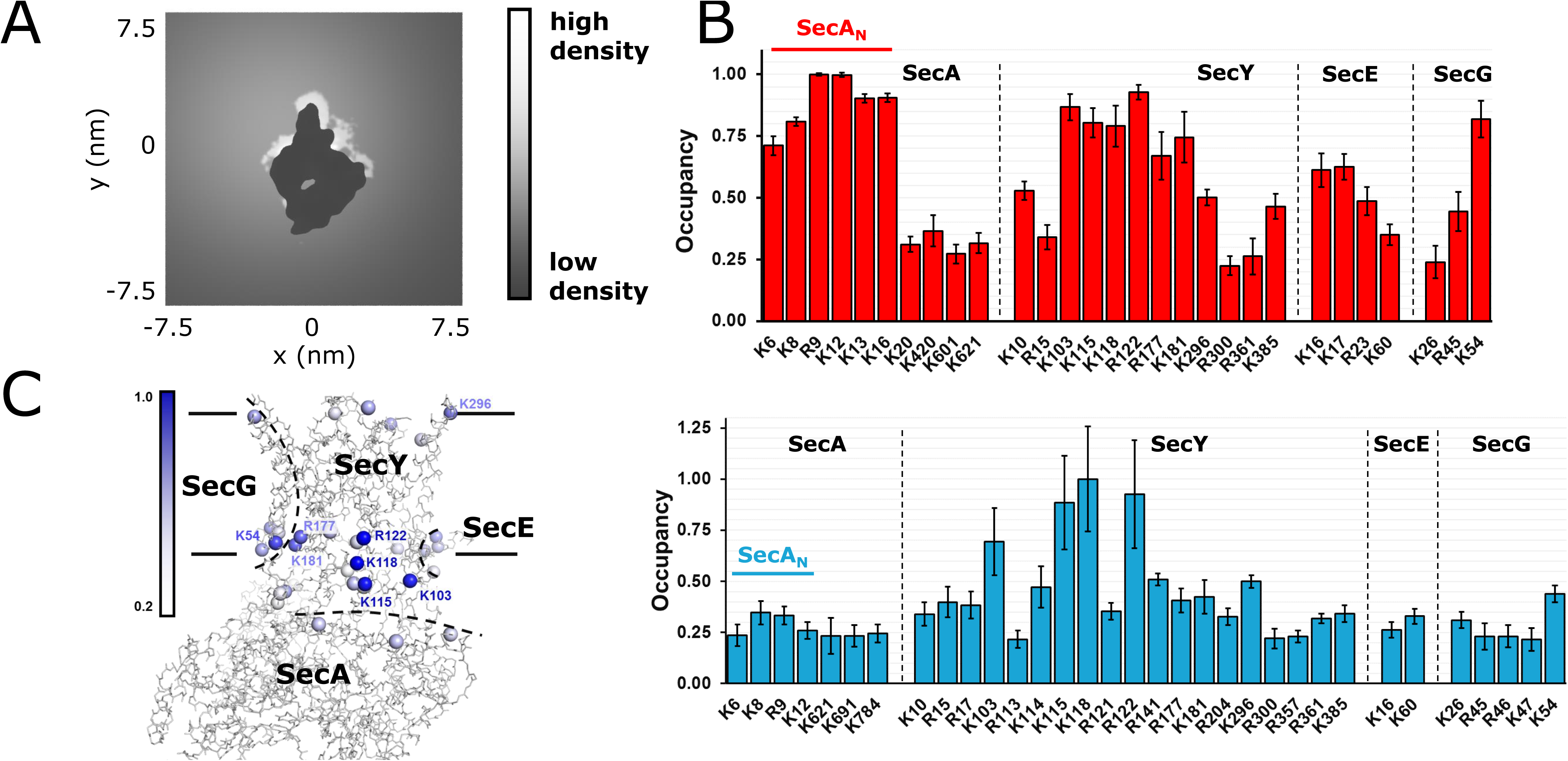
Distance analyses of Sec-CL. A) Density plot of CL head groups (‘PO1’, ‘PO2’ and ‘GL0’ beads; see Figure 1C) in the *xy* plane at the cytoplasmic face of the membrane. Data computed for the 10 simulations using the self-assembly method. The protein is shown in the centre of the plot in black. Distinct CL hotspots can be observed. B) Computed occupancies for CL binding to basic residues in each subunit of the Sec complex, with membranes formed using a self-assembly approach (red) and insane.py (blue). Data here have been normalised from 0-1, and error bars represent the s.e.m of 10 repeats (red) or five repeats (blue). All residues with less than 0.2 occupancy have been omitted for clarity. The full data set can be seen in Figure S1. SecA_N_, previously implicated in lipid binding (26), is indicated. C) The highest binding basic Sec residues are shown on SecA-SecYEG (PDB 3DIN (25)), with the CB atom of each residue coloured according to CL occupancy from the datasets in panel B, excluding SecA_N_. For clarity, only the residues with occupancy above 0.5 are labelled. The Sec subunits are approximately demarcated with black dotted lines. The position of the membrane is marked with black lines.

To probe these interactions in more detail, the distance between basic residues in SecA-SecYEG and nearby CL head groups was calculated using tools available in Gromacs (30) for each separate simulation. The data reveal multiple residues implicated in CL binding in SecA-SecYEG (Figure 2B; red data), including several residues with >90% binding occupancy (Figure S1; red data).

### Coarse-grained analyses of Sec CL interaction using preformed membranes

Next, we built five different mixed bilayers containing ~1% CL around SecA-SecYEG using the insane.py program (31). This lower concentration was chosen to reduce the high occupancy apparent in the self-assembly dataset (Figure S1; red data), thus allowing us to identify more specific Sec-CL interactions, and model the kinetics of CL binding and release. In addition, the restraints were removed from the protein beads to allow SecA-SecYEG to structurally react to the presence of the membrane, although additional harmonic bonds were added to maintain stability (see Methods). Each bilayer was simulated for an extended period of between 5 and 55 μs (sampling over 100 μs in total), with RMSD analyses of the protein BB beads confirming that the simulations were stable (Figure S2). As above, we determined the CL occupancy for each basic residue in the system, finding a range of 0-11% occupancy over the course of the simulation (Figure 2B; blue data).

The MD data highlighted several residues at which CL binding occurred. Those displaying the highest CL occupancy were K103, K115, K118, R122, R177 and K181 in SecY, and K54 in SecG (Figure 2C and Table 1). Of these, K103, K118, R122 and R177 in SecY are very poorly conserved across different bacterial species (Figure S3), and SecG is itself a dispensable subunit of the complex (3). Indeed, its removal does not affect anionic lipid interactions with the translocon (32). This led us to the conclusion that SecY residues K115 and K181 were of particular importance for CL binding.

In the case of CL bound to K181, additional contribution to binding appears to come from the acyl tails, as can be seen by the perturbed acyl tail order parameters (Figure S4). However, it should be noted that CL molecules in *T. maritima* likely have distinctive acyl tails, involving ether bonds between the glycerol and fatty acid and the presence of membrane-spanning acyl tails (33-35). As our simulations pair *T. maritima* SecA-SecYEG with *E. coli* CL, we were minded not to over-analyse the contribution to binding from the acyl tails.

In addition to these positions on SecY, we observe considerable CL binding to the 25 N-terminal residues of SecA (SecA_N_; Figure 2B & 3), which have previously been shown to be important for lipid binding (5, 26).

**Figure 3.**
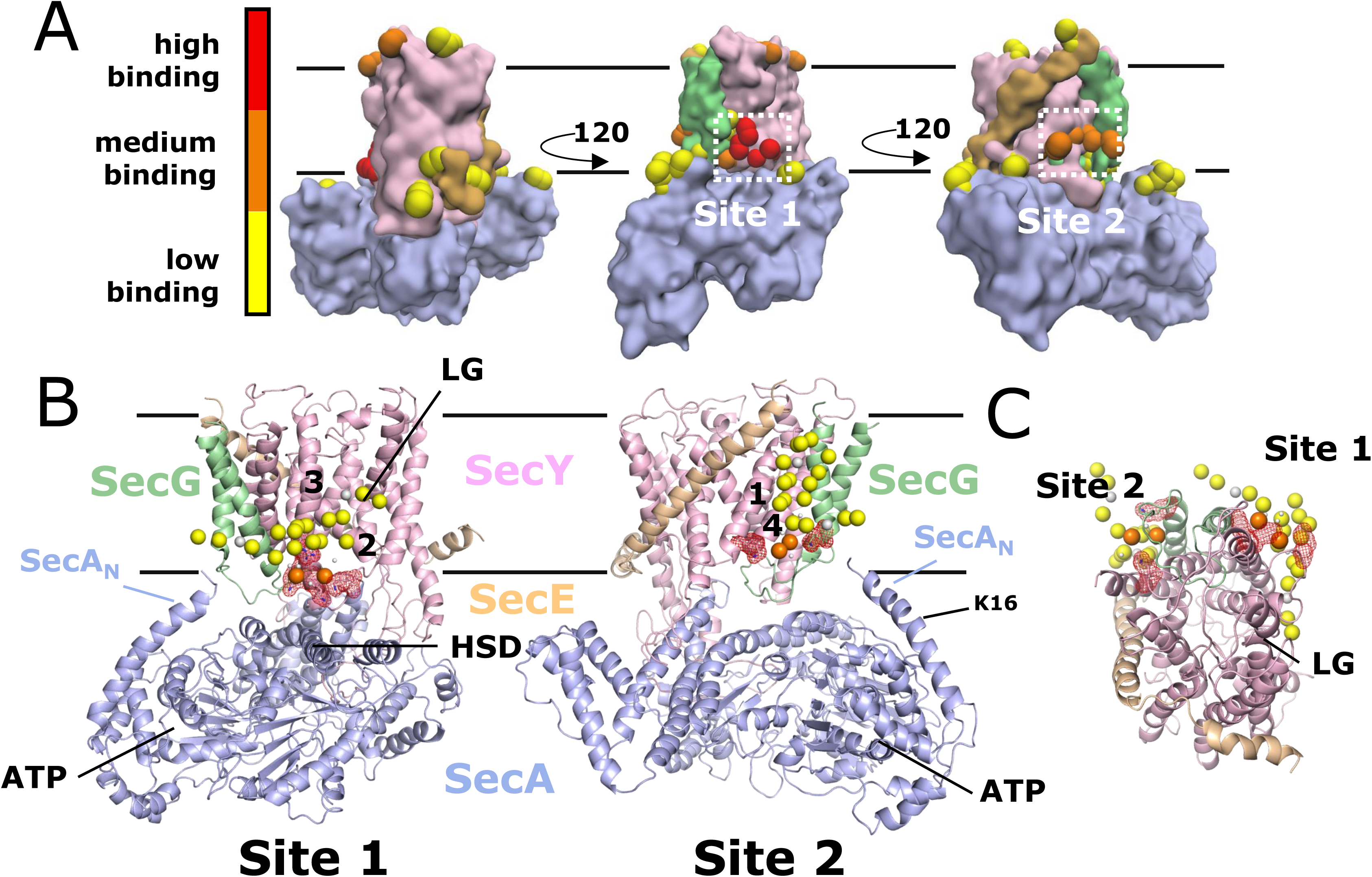
Structural views of CL binding. A) Views of SecA-SecYEG in MARTINI representation, shown as surfaces and coloured as per Figure 1A. CL-binding hotspots are shown as spheres and coloured according to normalised CL occupancy, with yellow beads representing occupancies of 0.2-0.5, orange 0.5-0.8 and red 0.8-1.0. Visual analysis reveals two distinct CL binding sites on the cytoplasmic face of SecY, labelled ‘Site 1’ and ‘Site 2’. B) Two frames taken from one of the CG simulation trajectories, with the protein atoms mapped back from CG to atomistic using the backward.py method (69). SecA-SecYEG is coloured as panel A. The CL molecule is shown in MARTINI representation, and coloured as per Figure 1C. Site 1 and 2 residues are shown as sticks and red mesh. Named SecY TMH are marked with numbers, with the SecY lateral gate (LG) and the SecA helical scaffold domain (HSD), N-terminus (SecA_N_; Lys-16 also labelled), and nucleotide binding site (ATP) labelled. C) As panel B, but with SecA removed and viewed from the cytoplasm. Both CL binding sites are shown. The molecule has been orientated as per Figure 2A.

### Kinetics of CL binding

Our longer (5-55 μs) simulations permit insight into the binding kinetics of CL to SecY. Distance analyses between K115/K181 and CL head groups reveals several clear binding events (Figure S5), where CL-residue distance is less than ~1 nm. A detailed analysis of the 55 μs simulation demonstrates that hundreds of binding events occur at both K115 and K181, lasting from ~100 ns to 2 μs, with a very broad sampling of binding times (Figure S6). Fitting the data to single exponentials reveals *k*_off_ values of ~4-5 μs^−1^ (Figure S6: red line). Kinetics are not always modelled accurately in the MARTINI force field (36), but these data clearly indicate that, at high concentrations of membrane CL (~ 10% = 70 mM) the on-rate would be very high indeed. Very fast binding and release of CL to the translocon, on the ns to μs timescale, is comparable with CL binding to other proteins (20, 21).

### Validation of CL binding sites by native mass spectrometry

Plotting a heat map of binding onto basic residues in *T. maritima* SecA-SecYEG reveals that the residues identified above form two distinct sites, termed here ‘Site 1’ and ‘Site 2’ (Figure 2B and Figure 3). Both sites are typified by the presence of multiple basic residues (Figure S7A). Site 1 comprises the higher CL-occupancy residues K103, K115, K118 and R122, which are all located on trans-membrane helices (TMHs) 2-3 of SecY, between the SecY lateral gate (LG), the SecA helical scaffold domain (HSD) and SecG (Figure 3B). Site 2 comprises K181 and R177 on SecY TMH 4, along with the mid CL-occupancy R15 and R17 on TMH 1. This site is close to the lipid-binding SecA_N_.

To validate the two CL binding sites, and to ascertain whether they are conserved across bacterial species, we designed *E. coli* SecYEG variants which abolish the primary positive charges: SecY^R113Q,R114Q,K115E,R121Q^EG for Site 1 and SecY^K20Q,R21Q,R22Q,R181E^EG for Site 2 (Figure S7B). We analysed the variants using native mass spectrometry (nMS (37-39)), observing peaks for SecE, SecG, SecEG, SecY, SecYG and SecYE (Figure S8A). The intact SecYEG complex was not detected due to the application of strong collisional activation energy in the mass spectrometer, required for detergent removal (in this case dodecyl maltoside; DDM) from the protein, which causes partial complex dissociation (40).

We see no CL binding to SecE, SecG, or SecEG, supporting the notion that CL binding occurs mainly to SecY. When using the Site 1 and Site 2 variants, we see a significant decrease in CL binding to SecY (Figure 4A), validating the assignment of the two CL binding sites from the CGMD.

**Figure 4.**
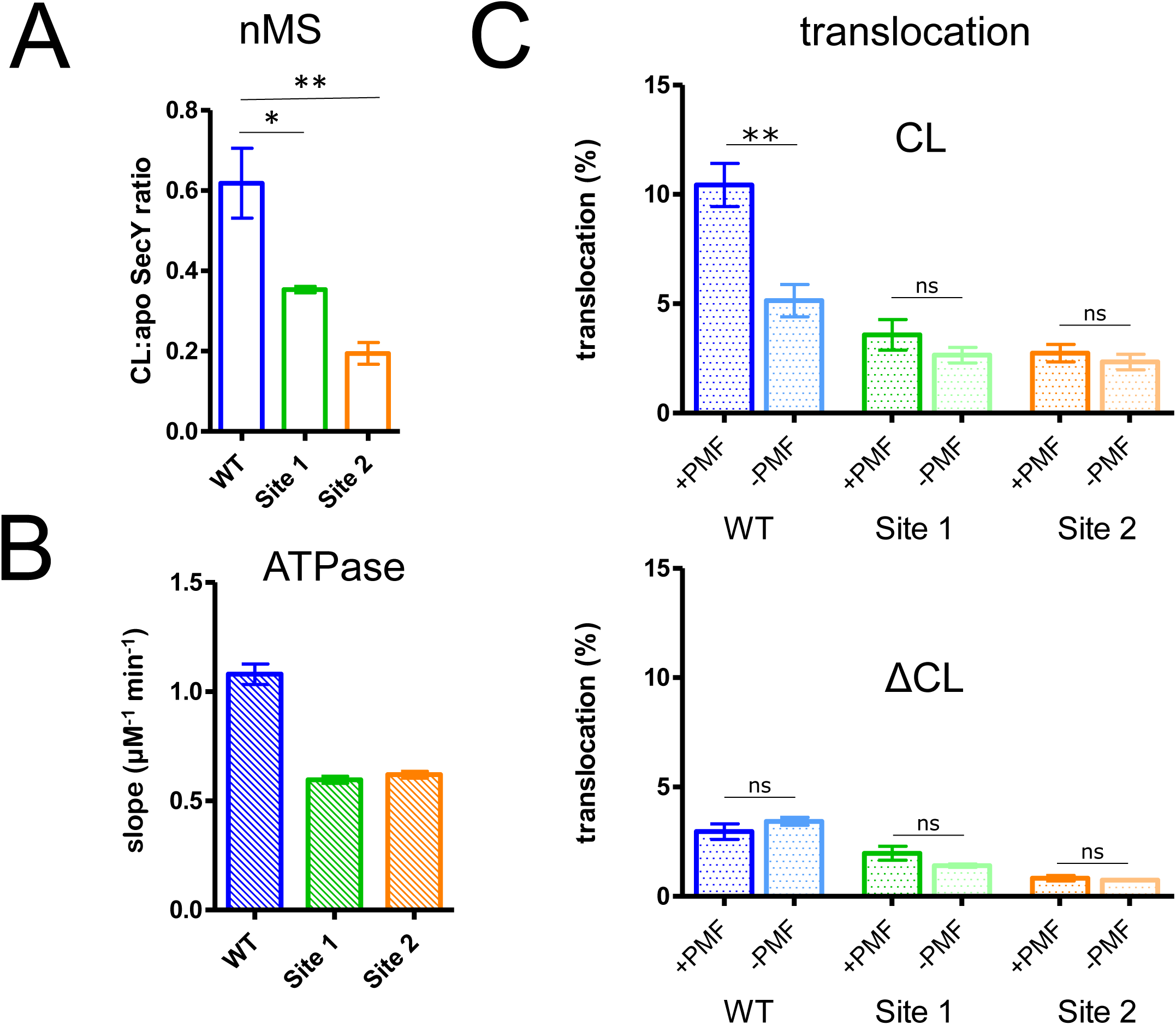
Experimental analyses of CL interaction. A) Ratios of CL-bound to non-bound (apo) SecY, as measured using nMS, where a value of 1.0 would be equal abundancies of both states. WT SecY has a significantly higher ratio of CL-bound SecY than either the Site 1 or Site 2 variants (*p* = 0.0194 and *p* = 0.0048 respectively, using a one-tailed *t*-test). For raw data see Table 3 and Figure S8A. B) ATPase analyses of SecA-SecYEG with increasing concentrations of CL. Shown are the rates of ATPase increase upon CL addition, as determined from fitting the titration data to linear slopes. Error bars are reported errors for the linear regression. See Figure S9A for titration data. C) *In vitro* translocation data for IMVs containing overexpressed WT or variant SecYEG. On the top are data for IMVs containing normal levels of CL (CL), which demonstrate a lower translocation efficiency in the Site 1 and Site 2 variants, and a total loss of PMF-stimulation. Below are data from IMVs with CL biosynthesis knocked out (ΔCL), for which none of the samples exhibit PMF stimulation. Error bars are s.e.m of 5 repeats, and reported statistical analyses are from one-tailed t-tests, where *p* = 0.0024, 0.261 and 0.462 for the CL data, and *p* = 0.302, 0.155 and 0.529 for the ΔCL data.

### Specific CL binding increases SecA-SecYEG ATPase activity

For each variant, we analysed the effect of CL on SecA stimulation. CL binding to SecYEG is known to stimulate SecA (10), so following SecA ATPase activity provides a readout of this process. We used CL titrations over a range of 0-40 μM, as CL appears to have an inhibitory effect on SecA ATPase activity above this concentration. When fitting the data to tight binding equations (Figure S9A, dotted lines) the inability to fit the full curve precludes accurate extraction of *K_d_* or *B*_max_. Therefore, to compare the sites qualitatively, we only fitted the linear concentration range (0-40 μM; Figure S9A, solid lines). The slopes from these fits reveal that CL stimulation of the Site 1 (0.60 ± 0.02 μM^−1^ min^-1^) and Site 2 (0.62 ± 0.02 μM^−1^ min^−1^) variants are reduced by almost half compared to wild-type SecYEG (WT; 1.08 ± 0.05 μM^−1^ min^−1^) (Figure 4B). The *k*_cat_ in the absence of CL is also elevated in the Site 1 variant (18.5 ± 0.31 min^−1^) relative to WT and Site 2 (8.6 ± 0.88 and 7.5 ± 0.30 min^−1^, respectively) (Figure S9A).

Between them, the sum of the two sites individually thus account for nearly 100% of CL binding to SecY, and for most of its stimulatory effect on ATPase activity. Therefore, these analyses strongly support the existence of two conserved CL-binding sites on SecY, predicted by CGMD.

### The role of specific CL binding in SecYEG function

Next, we addressed the physiological effects of CL binding at the two sites. We carried out translocation assays using inverted membrane vesicles (IMVs) over-expressing variant or WT SecYEG. The results revealed that both variants generally have a lower translocation efficiency, compared to WT SecYEG (Figure 4C & S9E, ‘CL’). Strikingly, whilst PMF-stimulation can be observed in the WT SecYEG IMVs, no PMF-stimulation can be observed in either variant. This strongly suggests a role for specific CL binding at Sites 1 and 2 in the stimulation of transport by the PMF.

To bolster these data, and ensure the observed lack of PMF stimulation was indeed due to loss of CL binding, we repeated the assay using IMVs from a strain of *E. coli* C43(DE3) in which all three CL biosynthesis pathways had been knocked out (27). We verified the knockdown of CL levels (Figure S9B-C) and confirmed the IMVs were able to produce a PMF (Figure S9D), *i.e.* CL is not essential for bacterial F-ATPase activity. Translocation data reveal that none of the IMVs exhibit PMF stimulation (Figure 4C & S9E, ‘ΔCL’), demonstrating that CL binding is required for the PMF stimulation of translocation.

The results signify that specific binding of CL to the sites identified by CGMD analyses is important to both ATP- and PMF-driven transport activity.

### Effect of CL binding on SecYEG dimerisation

To investigate the role of CL on SecYEG dimerization, further CGMD simulations were run on a system containing 16 copies of the *Thermus thermophilus* SecYEG translocon (41) in a membrane with either 0% or 10% CL. Visual analysis of the systems at 1.1 μs supports a role for CL in SecYEG dimerization (Figure 5A), although the dimeric forms observed are highly heterogenous (Figure 5A: SecE amphipathic helix shown in blue for context, as per Figure 5B). Following the number of protomers in a dimer or monomer reveals that after about 800 ns, there are consistently 4 dimers present in the CL-containing membrane (*i.e.* 8 monomeric complexes), but only 2 in the non-CL membrane (Figure 5C). The data suggest that CL has an important role in stabilising SecYEG dimerisation, as previously noted (10). However, whilst CL binding is observed at the equivalent Sites 1 and 2 (Figure 5D), in most cases this does not form part of the dimer binding interface (Figure 5D; top box), with any interfacial CL apparently interacting non-specifically (Figure 5D; bottom box). It is plausible that at least part of CL’s effect on driving SecYEG dimerization is through a more general mechanism, either stabilising each protomer or changing the biophysical properties of the membrane to encourage dimerisation. This stabilisation of an initial contact then might lead to re-arrangement to specific dimeric arrangements.

**Figure 5.**
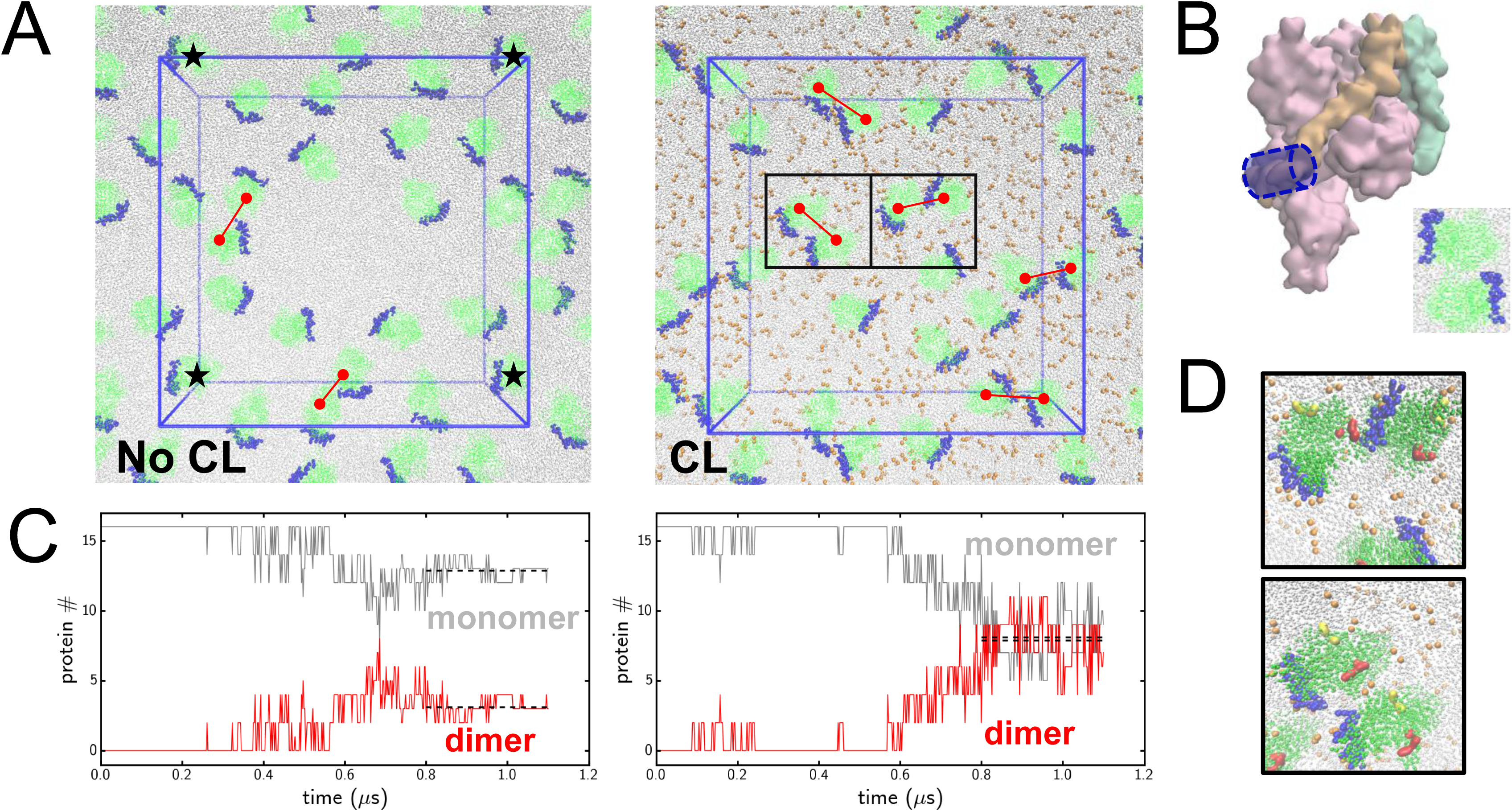
Role of CL in SecYEG dimerisation. A) Snapshots taken at 1.1 μs from simulations of 16 *T. thermophilus* SecYEG molecules (PDB 5AWW (41)) in membrane with either 0% or 10% CL. The membrane is shown in grey, with the CL head group phosphates as orange spheres, SecYEG beads are shown in green, except for the amphipathic helix of SecE in blue (as panel B). The periodic box is shown in blue, such that the SecYEG molecules marked with a black star are the same molecule. Dimers are shown with a red connecting line. Black boxes in the right image are the regions used in panel D. B) Molecule of SecYEG used in panel A showing the amphipathic helix of SecE. The most commonly accepted model of SecYEG dimerization involves the TMH of SecE interacting, meaning that the amphipathic helix of each protomer would be parallel but on opposite sides, as illustrated to the bottom right. C) Distance analyses were carried out between each SecYEG protomer and the nearest neighbouring SecYEG. The molecules were considered as dimers if the minimum distance to the next protomer was less than 0.9 nm. The formation of dimers was followed over time, revealing that after ~800 ns, 2 dimers are present without CL, whereas 4 dimers are present with CL. D) Close-ups from panel A. Here, Site 1 and 2 are shown as red and yellow surface respectively. In the top panel, CL binding can be seen at both Site 1 and 2, however CL appears to play no role in the dimer interface. In the bottom panel, CL is clearly acting as an interfacial lipid, however this is not mediated by binding to Sites 1 or 2.

## Discussion

Integral membrane proteins are strongly affected by the lipid environment in which they reside – through both specific and non-specific interactions (7, 8). Whilst structural data can occasionally reveal the binding of specific phospholipids to membrane proteins, more often the use of harsh detergents during purification precludes this. Thus, biophysical analyses of lipid binding can powerfully augment the structural data.

Here, we have employed the combined powers of computational (CGMD) and experimental (biochemical assays and mass spectrometry) approaches to identify and characterise specific CL binding sites on the bacterial Sec translocon.

The positions of the two identified CL-binding sites (termed here Site 1 and Site 2) are striking when considered in the context of SecA-SecYEG function. Site 1 is positioned on the edge of the lateral gate (LG), so that the bound CL contacts 2 of the 3 LG helices. Previous data have shown that the LG can adopt an open or closed state (2, 6, 25), with the transition between these two states implicated as an important feature of the protein translocation process (6). Thus, the heightened transport activity induced by specific binding of CL might relate to its effect on the equilibrium between these conformations. In addition, the LG acts as the site of signal sequence interaction with the membrane (42), previously shown to be important for regulating signal sequence structure (43, 44). It therefore seems plausible that there is a role for specific lipid interactions in productive signal sequence binding. Finally, CL binding at Site 1 will position the phosphate head group immediately adjacent to residues on the SecA helical scaffold domain (HSD) that have previously been shown to respond to lipid binding (45). This may be important in the activation of SecA, and likely explains the raised basal ATPase rate of the Site 1 variant (Figure S9A).

Site 2 is located on the other side of the channel, formed at the SecY-SecG interface. This site is in direct contact with the N-terminus of SecA, previously shown to be important for SecA-membrane binding in the *E.coli* system (5), particularly residues 23-25 (26). The CGMD data support this observation, with considerable CL binding observed for this region, including residue Lys-16 (Figure 3B “K16”), the *T. maritima* equivalent of the *E. coli* residue 23. A CL molecule bound to Site 2, therefore, might have an important role in SecA activation.

While SecYEG dimerization does not appear to be driven by specific interactions at Sites 1 or 2, this does not preclude their involvement in other protein-protein interactions. SecYEG can form a complex with the membrane ‘insertase’ YidC, along with the accessory subcomplexes SecDF and YajC, to form the ‘holo-translocon’, capable of protein secretion and membrane protein insertion (46, 47).The preliminary structure of the holo-translocon (48) predicts that Sites 1 and 2 are proximal to YidC in this complex; indeed, in this context the CGMD data predicts the positioning of the CL head group directly on the C1 region of YidC (Figure S10; (48, 49)). As with SecYEG alone, CL has been implicated in mediating the interaction of SecA with the HTL (46).

Together, these results demonstrate the importance of specific protein-lipid interactions to membrane protein structure and function, and the power of CGMD to identify and probe these interactions in other energy transducing membrane embedded machines. For the Sec machinery, we find that a specific and transient interaction of CL is critical for transport function. For many years we have known that by association CL is important for the respiratory and ATP synthase complexes. To our knowledge this is the first demonstration where a CL, or any lipid, has been directly implicated in the energy transducing process, rather than simply for complex stabilisation.

## Materials and methods

### Coarse-grained simulations

Simulations of SecA-SecYEG were built using the *T. maritima* crystal structure (PDB 3DIN (25)), with the nucleotide removed and missing loops remodelled in Modeller (50). Following this, two distinct setups were used, each described below.

Firstly, for the self-assembly method, atoms were converted to CG beads with the Martinize method (51), using MARTINI 2.2. While this approach has some limitations (36), it has been shown to be very powerful for the modelling of protein lipid interactions (52, 53). Next, the protein beads were built in a 15 nm cubic box, and 460 distearoyl phosphoethanolamine (DSPE) and 52 CL molecules were added before solvation and addition of counter ions. The CL parameters used are of bacterial di-1-palmitoyl-2-vaccinate-phosphatidylglycerol in a −2 charge state (54, 55). The systems were minimized using steepest descents for 2500 steps of 20 fs, before two rounds of equilibration: firstly 100 ps of NPT (constant-temperature, constant-pressure) equilibration at 300K with 2 fs time steps, then 5 ns of NPT equilibration at 340K with 20 fs time steps. Finally, simulations were run for 1 μs at 340K using 20 fs time steps, with positional restraints applied to the protein beads. The frames in which the bilayer had not yet formed – or in one case formed in the yz axis – were removed, and the simulations were analysed for CL density in VMD (Theoretical and Computational Biophysics group, University of Illinois at Urbana-Champaign), to identify possible CL interaction sites.

Alternatively, for the insane.py method, symmetrical membranes of 9 CL and 823 dipalmitoyl phosphatidylcholine (DPPC) molecules were built around the protein beads using insane.py (31), modified here for the inclusion of CL. The protein beads were described using MARTINI 2.2, with additional harmonic bonds applied to supplement the non-bonded interactions and help preserve the higher-order structure of the input models, with omission of these bonds leading to considerable deformation of the protein. The bonds had a force constant of 500 kJ mol^−1^ nm^−2^ and an upper bond length cut-off of 0.9 nm. Next, the systems were minimized and equilibrated as described above, before production simulation. For these simulations, RMSD analyses were carried to ensure stability of the protein beads (Figure S2). These simulations were largely run on EPCC‘s Cirrus HPC Service and the ARCHER UK National Supercomputing Service.

As well as modelling of the SecA-SecYEG systems, models were built from *T. thermophilus* SecYEG molecule (PDB 5AWW (41)). For this, 16 protomers were aligned evenly across a 40 × 40 × 12 nm box, and built into a membrane of either 3499 DPPC and 388 CL molecules (“CL”) or 3888 DPPC molecules with no CL (“NoCL”) using the insane.py method (31). The systems were minimized and equilibrated as described above, before further equilibration for 200 ns and finally a 1.1 μs production simulation. Dimer formation was carried out through iterative minimum distance analyses for each protomer, where a distance of >0.9 nm was considered to be bound in a dimer.

All simulations were run in Gromacs 5.1.2 (30). Non-bonded interactions were treated with a switch function from 0-1.2 nm and 0.9-1.2 nm for the Coulomb and Lennard-Jones interactions respectively. Temperature coupling was achieved with the Bussi-Donadio-Parrinello thermostat (56), and semi-isotropic pressure coupling was by the Parrinello-Rahman barostat (57, 58).

Images of proteins were made in PyMOL (59) or VMD (29). Graphs were plotted using Microsoft Excel, GraphPad Prism or Matplotlib.

### Protein expression and purification

Mutations were introduced using the QuikChange protocol (Stratagene) to *secEYG* in a pBAD expression plasmid. The SecYEG variants were expressed as previously described (60); briefly, each variant was expressed in C43(DE3) *E. coli,* which were lysed and the membranes isolated by ultracentrifugation. The SecYEG protein was solubilised from the membrane fraction using 1 % DDM (n-Dodecyl β-D-maltoside) and purified using nickel chromatography (in 0.5 % C12E9 for the CL-titration experiments or in 0.1 % DDM for the nMS) followed by size exclusion chromatography and reverse ion-exchange chromatography (in 0.1 % C12E9 for the CL-titration experiments or in 0.02 % DDM for the nMS). The C12E9 purification has previously been shown to remove all bound CL, while the DDM purification allows some CL to remain bound (10). The purified proteins were concentrated and aliquoted for storage at −80°C.

SecA and the model substrate proOmpA were expressed and purified as described previously (61). To facilitate quantification by western blot, a C-terminal minimal V5 tag (IPNPLLGL) was added to the proOmpA gene by PCR, and confirmed by DNA sequencing.

### ATPase assays

*In vitro* ATPase assays were performed as described previously (61, 62), with 0.3 μM SecA and 2.4 μM SecYEG in 0.1 % C12E9, in TK_150_M buffer (20 mM Tris pH 7.5, 150 mM KCl, 2 mM MgCl_2_) with 0.03 % C12E9. Reactions were followed by coupling ATP hydrolysis to NADH oxidation, measuring absorbance at 340 nm. To each reaction, a set amount of CL in 0. 6 % C12E9 was added to produce a final concentration of between 0 and 80 μM. Note that above 40 μM, CL appears to have an inhibitory effect on SecA ATPase. Fixed volumes of each reagent were used to ensure that identical C12E9 concentrations were present in each reaction (0.1 %). Data were initially fitted to a tight-binding equation (Figure 9A; dotted line (6)), but the quality of fit was too low to extract meaningful information from. Data from 0-40 μM were therefore fitted by linear regression instead.

### Native mass spectrometry

Capillaries for nMS were produced using a Model P-97 capillary puller (Sutter Instruments) and gold coated using a Q150RS sputter coater (Quorum). Purified SecYEG in 0.02 % DDM was exchanged into MS suitable buffer (250 mM ethylenediamine diacetate (EDDA), 0.014 % DDM (v/v)) to a final concentration of 9 μM (SecYEG WT) or 12μM (SecYEG variants) using Micro Bio-Spin 6 columns (Bio-Rad). 1.5 μL protein sample was then loaded into the gold-coated capillaries and introduced into a Synapt G2-Si (Waters) mass spectrometer by nano-electrospray ionisation. The following conditions were used in the mass spectrometer: capillary voltage +1.2 kV, sampling cone voltage 20-50 V, backing pressure 3.88 mbar, trap and transfer pressure (argon) 1.72 e^−2^ mbar, ion mobility cell pressure (nitrogen) 2.58 mbar. Trap and transfer cell collision energies (CE) were adjusted to remove the DDM detergent micelle, allowing the spectra to be resolved. Spectra were recorded at trap CE 200V and transfer CE 200V. Mass measurements were calibrated using caesium iodide (100mg/ml). Spectra were recorded and smoothed using Masslynx 4.1 (Waters) software.

The relative abundances of each oligomeric/lipid bound state of SecYEG was calculated with a Bayesian deconvolution software, UniDec (63). Detector efficiency was accounted for before quantification (64). The spectra were analysed between 3,000-8,000 m/z to allow the quantification of the SecY based complexes. SecE and SecG monomers were found in the 1,000-3,000 m/z range and were discounted from our analysis as they were detected in vast excess to the other oligomeric species. The mass range for peak detection was 45,000-77,000 Da and species were identified under the following masses: SecY 48,325 ± 50 Da; SecY + CL 49,700 ± 50 Da; SecYG 59,725 ± 50 Da; SecYG + CL 61,125 ± 50 Da, SecYE 63,050 ± 50 Da; SecYE + CL 64,425 ± 50 Da; SecYEG 74,550 ± 50 Da. The comparison of the apo:CL bound relative abundances assumes lipid bound species have similar ionisation efficiencies.

We see CL binding to SecY, SecYG and SecYE. For the SecYG and SecYE peaks, no significant difference is seen (Figure S8B) between the WT and variant samples. This indicates that the presence of SecE and SecG are able to compensate for the loss of the binding sites on SecY in our Site 1 and 2 variants. This is consistent with the observed higher quantities of CL associated with SecYG and SecYE, compared to SecY alone (Tables 3-4). Part of this effect is most likely through a contribution of SecE and SecG to the CL binding sites – particularly through provision of a hydrophobic subunit interfaces for the acyl tails, *e.g.* as seen between SecY and SecG (Figure 3B). In addition, part of the effect may come through a stabilisation of the correct SecY conformation, which would explain the higher abundancies of SecYE and SecYG, compared to SecY alone (Table 3).

### Preparation of inverted membrane vesicles

Inverted membrane vesicles (IMVs) were made from C43(DE3) *E. coli* cells, as well as a C43(DE3) strain in which the three CL synthase genes (*clsA*, *clsB* and *clsC*) have been knocked out (27). All cells were grown in a modified form of M9 minimal media (12.8 g Na_2_HPO_4_, 3 g KH_2_PO4 0.5 g NaCl 1 g NH_4_Cl, 20 ml glycerol, 2 mM MgSO_4_, 100 μM CaCl_2_ and 10 μM FeSO_4_). The cells were lysed, and centrifuged at 18,000 g for 20 mins to clear the lysate, followed by centrifugation on a bed of 20% sucrose at 110,000 g for 2 hours. The pellets were resuspended in TK_150_M, homogenised and run on a stepped sucrose gradient of 0–1.6 M sucrose for ~16 hours at 165,000 g. The IMV band was detected visually, extracted and diluted in TK_150_M, and centrifuged 350,000 g for 90 minutes. The pellets were resuspended in TK_150_M and aliquoted for storage at −80°C.

### Lipid quantification mass spectrometry

Both sets of cells were grown in M9 minimal medium until OD ~0.8 before induction of WT SecYEG. 500 ml of cells were harvested at 4000 g, resuspended in 50 ml of 20 mM Tris pH 7.5 with 300 mM NaCl, and then finally centrifuged at 2500 g. After weight estimation, pellets were resuspended in MilliQ water at a 1:2 mass/volume ratio. Samples were sonicated for 30 min at 30% power using a 50% pulse. After lipid extraction, as described in (65), phospholipid analysis was performed using liquid chromatography-electrospray ionization-tandem mass spectrometry (LC-ESI-MS) or HPLC (RSLC Dionex-U3000) equipped with a Corona-CAD Ultra detector coupled to a LTQ-Orbitrap Velos Pro mass spectrometer according to (66), and supplied by ThermoFisher Scientific.

Note that the C43(DE3) cells were diluted 20 fold for the analysis, and quantification was achieved from the CAD chromatogram. The ΔCL cells were diluted 3 fold, and approximate quantification achieved by extraction of the ionic current in the mass windows corresponding to CL.

### Translocation assays

Transport assays were performed using a modified version of the standard protocol (67). IMVs at a concentration of 0.4 μM SecY were mixed with 1 μM SecA, 0.2 μM proOmpA and an ATP regenerating system (0.1 mg ml^−1^ creatine kinase and 5 mM creatine phosphate) in TK_150_M. Under these conditions, ATP both powers translocation and generates a PMF *via* F-ATPase.

Reactions were prewarmed to 25°C then started by the addition of 1 mM ATP. After 200 seconds, reactions were quenched by the addition of the same volume of proteinase K at 0.6 mg/ml in 50 mM HEPES pH 8, and incubated on ice for 30 mins. The proteinase K step ensures that any untranslocated material will be digested. The samples were then precipitated with TCA added to 20%, before centrifugation for 10 minutes at 10,000 g. Pellets were dried and resuspended in 4X LDS gel loading buffer, before analysis with SDS-PAGE and western blotting against the V5 tag using a V5 tag antibody (SV5-Pk1, GeneTex) followed by a DyLight 800 anti-mouse secondary (SA5-10172, ThermoFisher). Gels were imaged in a LI-COR Odyssey, and bands were quantified in Image Studio.

## Acknowledgements

This work was funded by the BBSRC (BB/M003604/1, BB/I008675/1 and BB/N015126/1), and the Wellcome Trust (104632 and 109854/Z/15/Z). The MS work was supported by the Centre National de la Recherche Scientifique, INSERM, the “Initiative d‘Excellence” program from the French State (Grant “DYNAMO”, ANR-11-LABEX-0011-01) (to MC) and the Région Ile de France for co-funding the SAMM Mass spectrometry Facility at IPSIT (Chatenay-Malabry, France).EP is the recipient of an Imperial College London Institute of Chemical Biology EPSRC CDT studentship. This work was carried out using the computational facilities of the Advanced Computing Research Centre, University of Bristol (http://www.bris.ac.uk/acrc/). Additional simulations were carried out using computer time on EPCC’s Cirrus HPC Service (https://www.epcc.ed.ac.uk/cirrus) and on the ARCHER UK National Supercomputing Service (http://www.archer.ac.uk), provided by HECBioSim, the UK High End Computing Consortium for Biomolecular Simulation (hecbiosim.ac.uk), supported by the EPSRC. The authors thank Audrey Solgadi for MS analysis at IPSIT.

